# Comparative genomics of sex determination related genes reveals shared evolutionary patterns between bivalves and mammals, but not *Drosophila*

**DOI:** 10.1101/2025.01.30.635362

**Authors:** Filippo Nicolini, Sergey Nuzhdin, Fabrizio Ghiselli, Andrea Luchetti, Liliana Milani

## Abstract

The molecular basis of sex determination (SD), while being extensively studied in model organisms, remains poorly understood in many animal groups. Bivalves, a diverse class of molluscs with a variety of reproductive modes, represent an ideal yet challenging clade for investigating SD and the evolution of sexual systems. However, the absence of a comprehensive framework has limited progress in this field, particularly regarding the study of sex-determination related genes (SRGs). In this study, we performed a genome-wide sequence evolutionary analysis of the Dmrt, Sox, and Fox gene families in more than 40 bivalve species. For the first time, we provide an extensive and phylogenetic-aware dataset of these SRGs, and we find support to the hypothesis that *Dmrt-1L* and *Sox-H* may act as primary sex-determining genes, by showing their high levels of sequence diversity within the bivalve genomic context. To validate our findings, we studied the same gene families in two well-characterized systems, mammals and *Drosophila*. In the former, we found that the male sex-determining gene *Sry* exhibits a pattern of amino acid sequence diversity similar to that of *Dmrt-1L* and *Sox-H* in bivalves, consistent with its role of master SD regulator. In contrast, no such pattern was observed among genes of the fruit fly SD cascade, which is controlled by a chromosomic mechanism. Overall, our findings highlight similarities in the sequence evolution of some mammal and bivalve SRGs, possibly driven by a comparable architecture of SD cascades. This work underscores once again the importance of employing a comparative approach when investigating understudied and non-model systems.

## INTRODUCTION

In sexually reproducing organisms, the mechanism of sex determination (SD), i.e., the process by which the male or female identity of an organism (or gonadic tissue) is established, is highly diverse, ranging from strictly genetic systems to environmentally-dependent processes (**Haag & Doty, 2005**; **Uller & Helanterä, 2011**; **Bachtrog et al., 2014**; **Beukeboom & Perrin, 2014**). Characterising the molecular basis of SD is crucial for understanding not only the reproductive biology but also the evolutionary pressures shaping these systems (**Wilkins et al., 1995**; **Ellegren & Parsch, 2007**; **Grath & Parsch, 2016**; **Nicolini et al., 2023a**), as sex-determination related genes (SRGs; including primary sex-determining genes [SDGs]) are those responsible for the phenotypic differences between males and females, thanks to their sex-biased expression and interactions (**Ellegren & Parsch, 2007**; **Beukeboom & Perrin, 2014**; **Grath & Parsch, 2016**). One key aspect of SRGs is that they often exhibit accelerated rates of sequence evolution, due to their involvement in sex-related traits and reproduction. This represents the effect of sexual selection and/or adaptation, which act on sex-biased genes and produce highly divergent proteins at the interspecific level (**Civetta & Singh, 1998**; **Ellegren & Parsch, 2007**; **Meisel, 2011**; **Grath & Parsch, 2016**). Rapid sequence evolution is known for *Sex-determining Region Y* (*Sry*) of therians (**Pamilo & O’Neill, 1997**; **Mawaribuchi et al. 2012**), *Doublesex and mab-3 related gene W* (*Dm-W*) of the African clawed frog *Xenopus laevis*, and *Doublesex and mab-3 related gene Y* (*Dmy*) of the medaka fish *Oryzias latipes* (**Mawaribuchi et al. 2012**), all of which are master SDGs, that is, genes whose expression is primarily responsible for the establishment of the sex of the organism. Evolution under episodic diversifying selection has been detected also in *Drosophila* for genes involved in the SD cascade (e.g., *Sex-lethal* [*Sxl*], *transformer* [*tra*], and *doublesex* [*dsx*]), in correspondence with its establishment in the genus common ancestor (**Mullon et al., 2012**; **Baral et al., 2019**); though, rapid sequence evolution does not seem to concern extant amino acid sequences (**Haerty et al., 2007**; **Baral et al., 2019**), as they are globally evolving under purifying selection, especially in their catalytic domain (**Mullon et al., 2012**; **Baral et al., 2019**). However, concerning the *dsx* genes, higher rates of nucleotide and amino acid sequence evolution can be observed for male-specific regions, if compared to female-specific regions and dimerization domains (**Baral et al., 2019**).

While SD has been extensively studied in model organisms, like mammals, insects, and roundworms, comparatively little is known about the molecular mechanisms in non-model organisms. A remarkable example is represented by bivalve molluscs, which exhibit a wide variety of reproductive modes and sexual systems (**Breton et al., 2018**). Notwithstanding the considerable importance in the human socio-economic landscape (reviewed in **Haszprunar & Wanninger, 2012**; **Gomes-dos-Santos et al., 2020**), the study of SD mechanisms in bivalves has been hampered by the striking divergence among species (**Li et al., 2022**), and thus largely overlooked and limited to few case studies (**Breton et al., 2018**; **Nicolini et al., 2023a**). So far, no master SDG has been unambiguously identified, and the only working hypothesis on the functioning of the SD gene regulatory network is available for the Pacific oyster *Magallana* (formerly *Crassostrea*) *gigas* (**Zhang et al., 2014**). Nonetheless, the field still lacks both a robust functional investigation and an evolutionary framework in which to place the current knowledge (**Nicolini et al., 2023a**). As a matter of fact, major efforts have been dedicated to identify sex-biased genes through differential gene expression (DGE) analyses (e.g., **Ghiselli et al., 2012**; **Milani et al., 2013**; **Teaniniuraitemoana et al., 2014**; **Zhang et al., 2014**; **Capt et al., 2018**; **Afonso et al., 2019**), but very few have leveraged cutting-edge techniques to investigate their actual role in SD and/or gonad differentiation and development (e.g., **Liang et al., 2019**; **Sun et al., 2022**).

Components of the *doublesex* and *mab-3* related transcription factor (Dmrt), *Sry*-related HMG-box (Sox), and forkhead box (Fox) gene families (henceforth referred to as DSFG) are notoriously known as key actors in several developmental processes across Metazoa (**Benayoun et al., 2011**; **Matson & Zarkower, 2012**; **Sarkar & Hochedlinger, 2013**; **Mawaribuchi et al., 2019**), including SD in certain clades: the aforementioned *Dm-W*, *Dmy*, and *dsx* all belong to the Dmrt gene family, while *Sry* belongs to the Sox gene family; *Fox-L2*, which takes part in most of the vertebrate SD processes as a downstream effector of the female pathway, belongs to the Fox gene families. Members of the DSFGs have been identified as putative SRGs also in bivalves, thanks to both DGE analyses and *in-situ* hybridization (ISH; e.g., **Naimi et al., 2009**; **Li et al., 2018**; **Liang et al., 2019**; **Yue et al., 2021**), suggesting that their role in morphological and sexual development is maintained also in the clade. However, the clear role of DSFGs has yet to be elucidated, probably as a consequence to the lack of (i) a systematic classification of the gene families and (ii) a comprehensive understanding of their evolutionary history.

To overcome such limitations, this study aims to perform a thorough investigation of the DSFG families in bivalves, with the attempt to provide a high-quality resource to be used as a reference for future studies. Through the analysis of more than 40 annotated bivalve genomes and transcriptomes, we aim (i) to describe the complete set and evolutionary history of DSFGs in bivalves by means of phylogenetic inferences, manual curation, and orthology prediction; (ii) to identify DSFGs potentially involved in bivalve SD by investigating their sequence evolution in a genome-wide context. Our hypothesis is that, if any of the DSFGs is directly involved in SD (i.e., is a SDG), then we should expect it to show a higher rate of sequence evolution, as already found in previous studies (**Pamilo & O’Neill, 1997**; **Mawaribuchi et al. 2012**) and discussed earlier; this characteristic, in turn, would be reflected in a high diversity of the extant amino acid sequences across the bivalve clade. To assess the robustness and reliability of our approach, we additionally applied our pipeline to two non-bivalve datasets, composed of mammal and *Drosophila* species, respectively (hereon referred to as the ‘mammal dataset’ and the ‘fruit fly dataset’). By choosing two clades for which SD is well characterised, we wanted to compare our results with those obtained on taxa for which a deeper and more detailed knowledge is available. In particular, mammals and *Drosophila* provide two different frameworks to study the patterns of molecular evolution in SDGs: the former is a system where SD is completely genetic (i.e., the development into a male or into a female is triggered by the up- or downregulation of *Sry* in undifferentiated gonads, respectively), and for which a high rate of sequence evolution is already known for the male SDG *Sry*; the latter is instead a system where SD is chromosomic, thus lacks a master SDG (the sexual fate of the individual is determined by the ratio between autosomes and X chromosomes), and no high rate of sequence evolution has been detected in extant amino acid sequences of SRGs (which are globally evolving under purifying selection). In this sense, the mammal and the fruit fly datasets represent opposing control datasets to be compared to bivalves, as it is expected that a higher rate of sequence evolution concerns only master SDGs (like *Sry* in therians; i.e., the top regulatory part of the SD cascade), but not the downstream genes (i.e., the bottom effectors). Therefore, we test our pipeline on mammals and fruit flies with the following expectations: (i) in the mammalian dataset, *Sry* should be detected as fast-evolving (**Pamilo & O’Neill, 1997**; **Mawaribuchi et al. 2012**); while (ii) in the fruit fly dataset, no gene among those working within the sex-determining cascade should result as evolving at a higher pace (**Haerty et al., 2007**; **Mullon et al., 2012**; **Baral et al., 2019**). In this way, we are able to assess the reliability of results in bivalves, by comparing them to a null hypothesis.

This work offers novel insights into the evolutionary dynamics of SRGs and contributes a valuable genomic resource for understanding SD in bivalves, one of the most ecologically and economically important groups of marine organisms. In particular, here we provide the first extensive phylogenetic-based classification of DSFGs in bivalves, covering many species from the major bivalve orders, along with a comprehensive investigation of their sequence evolution.

## MATERIALS AND METHODS

### Dataset of bivalve annotated genomes and transcriptomes

Annotated genome assemblies of bivalves, gastropods, and cephalopods were obtained from various publicly available resources (**Supp. Tab. S1**). Isoforms from genome annotations were removed using the AGAT toolkit (v0.8.0; **Dainat, unpublished**). The nucleotide coding sequences for *Sinonovacula constricta* (Adapedonta) was not available for download. To avoid excluding the species from our analyses, the corresponding file was generated in-house by mapping the annotated protein sequences on the reference genome using miniprot (v0.13-0; **Li, 2023**). In order to provide an extensive identification of SRGs also for underrepresented bivalve orders (mainly belonging to the Heterodonta clade), 14 additional species represented by sequenced transcriptomes were included in the analyses, and were obtained from **Piccinini et al., 2021**, **Iannello et al., 2023** and **Iannello et al., in preparation**. The resulting set of annotated genomes and transcriptomes (hereafter referred to as the ‘comprehensive bivalve dataset’) was checked for completeness using BUSCO with the Metazoa reference dataset (v5.2.2; **Manni et al., 2021**).

### Identification and classification of Dmrt, Sox and Fox genes in bivalves

Members of DSFG families were retrieved from the comprehensive bivalve dataset with hmmsearch (v3.3.2; http://hmmer.org/). The signature catalytic domains of the DSFG families were used as queries. Specifically, HMM profiles were built after the Pfam databases for the DM domain (PF00751), the HMG box (PF00505), and the forkhead domain (PF00250), respectively. Obtained hits were then annotated using (i) the PANTHER HMM standalone sequence scoring against the PANTHER library v18.0 and (ii) RPS-BLAST (v2.5.0+) against CDD (pre-compiled version, downloaded on 09/11/23). In both cases, hits with an E-value of 10^−5^ were retained. Genes which were correctly annotated by both systems (**Supp. Tab. S2**) were kept for subsequent analyses. DSFGs from *Homo sapiens*, *Drosophila melanogaster*, and *Caenorhabditis elegans* (**Supp. Tab. S3**; the reference species) were retrieved from NCBI and were used as reference genes for annotation (see below). Classification and nomenclature of each family were retrieved from: **Mawaribuchi et al., 2019** for Dmrt genes; **Phochanukul & Russell, 2010** and **Sarkar & Hochedlinger, 2013** for Sox genes; **Mazet et al., 2003** for Fox genes. The alignments of mollusc and reference DSFGs were guided by the aforementioned Pfam HMM profiles and performed with Clustal Omega (v1.2.3; **Sievers et al., 2011**), then trimmed with trimAl (v1.4.rev15; **Capella-Gutiérrez et al., 2009**) with a gap threshold of 40%. Resulting alignments were manually inspected to remove sequences with incomplete catalytic domains, then aligned and trimmed again as before. Phylogenetic trees were inferred using IQ-TREE (v2.1.4-beta COVID-edition; **Minh et al., 2020**) with automatic model selection (**Kalyaanamoorthy et al., 2017**) and 1,000 bootstrap replicates. The phylogenetic tree of Dmrt genes was midpoint rooted, as no clear outgroup has been found so far (**Wexler et al., 2014**). Phylogenetic trees of Sox and Fox gene families were rooted using two fungi MATA-1 sequences (XP_62685912.1, CCD57795.1) and two Amoebozoa forkhead-like domains (XP_004368148.1, XP_004333268.1), respectively (**Heenan et al., 2016**; **Nakagawa et al., 2016**). The rooting was performed with Gotree (v0.4.5; **Lemoine & Gascuel, 2021**). To identify and annotate bivalve homology groups within each gene family, we utilized the algorithm implemented in Possvm (v1.2; **Grau-Bové & Sebé-Pedrós, 2021**), with DSFGs from the reference species as reference annotation. To better establish the orthology relationships among ambiguous groups of Dmrt and Fox genes, we ran a series of other phylogenetic reconstructions (see **Discussion**), by using the same pipeline as before. In the case of *Fox-Y* genes, we also used Fox gene sequences from the sea urchin *Strongylocentrotus purpuratus*, as given by **Tu et al. (2006)**. All the phylogenetic trees were plotted using the R package ‘ggtree’ (**Yu et al., 2017a**).

### Sequence diversity of bivalve single-copy orthogroups

As a metrics to measure the sequence diversity of bivalve DSFGs, and test whether those putatively involved in SD showed higher values than other genes, we used the amino acid sequence divergence (AASD). This metric is fast and straightforward to obtain, as it only requires the amino acid alignment and the corresponding best-fit substitution model. To this purpose, we produced amino acid alignments of bivalve single-copy orthologous (SCOs) groups and built the distribution of their median AASD. To further save computational time and prevent over-represented bivalve clades (such as Ostreida and Mytilia) to bias the analysis, SCOs were obtained after a reduced dataset comprised of 34 bivalve species (hereafter referred to as the ‘reduced bivalve dataset’; **Fig. 1**; **Supp. Tab. S1**), which included, for each bivalve genus, only the best genomes and transcriptomes in terms of either BUSCO scores or assembly statistics (**Supp. Tab. S1**). *Archivesica marissinica* and *Saccostrea glomerata* were removed, as their annotated coding sequences contain many stop codons. Genes were clustered in orthogroups using OrthoFinder (v2.5.5; **Emms & Kelly, 2019**) with DIAMOND ultra-sensitive and default parameters. Resulting orthogroups were splitted into SCOs composed by at least 17 species (50% of the bivalve reduced dataset) using DISCO (v1.3.1; **Willson et al., 2022**). Amino acid and nucleotide sequences were then aligned using Clustal Omega as implemented in TranslatorX (v1.1; **Abascal et al., 2010**), and jointly trimmed using trimAl with a gap threshold of 40% and the removal of spurious sequences (-resoerlap 50 -seqoverlap 50). Eventually, SCOs containing (i) internal stop codons, (ii) with less than 17 species left, or (iii) containing DSFGs were removed from downstream analyses. The best amino acid substitution model was inferred for each trimmed alignment using ModelFinder as implemented in IQTREE2 (model search was restricted to the following matrices: Blosum62, cpREV, Dayhoff, DCMut, FLU, HIVb, HIVw, JTT, JTTDCMut, LG, mtART, mtMAM, mtREV,

**Fig. 1.**
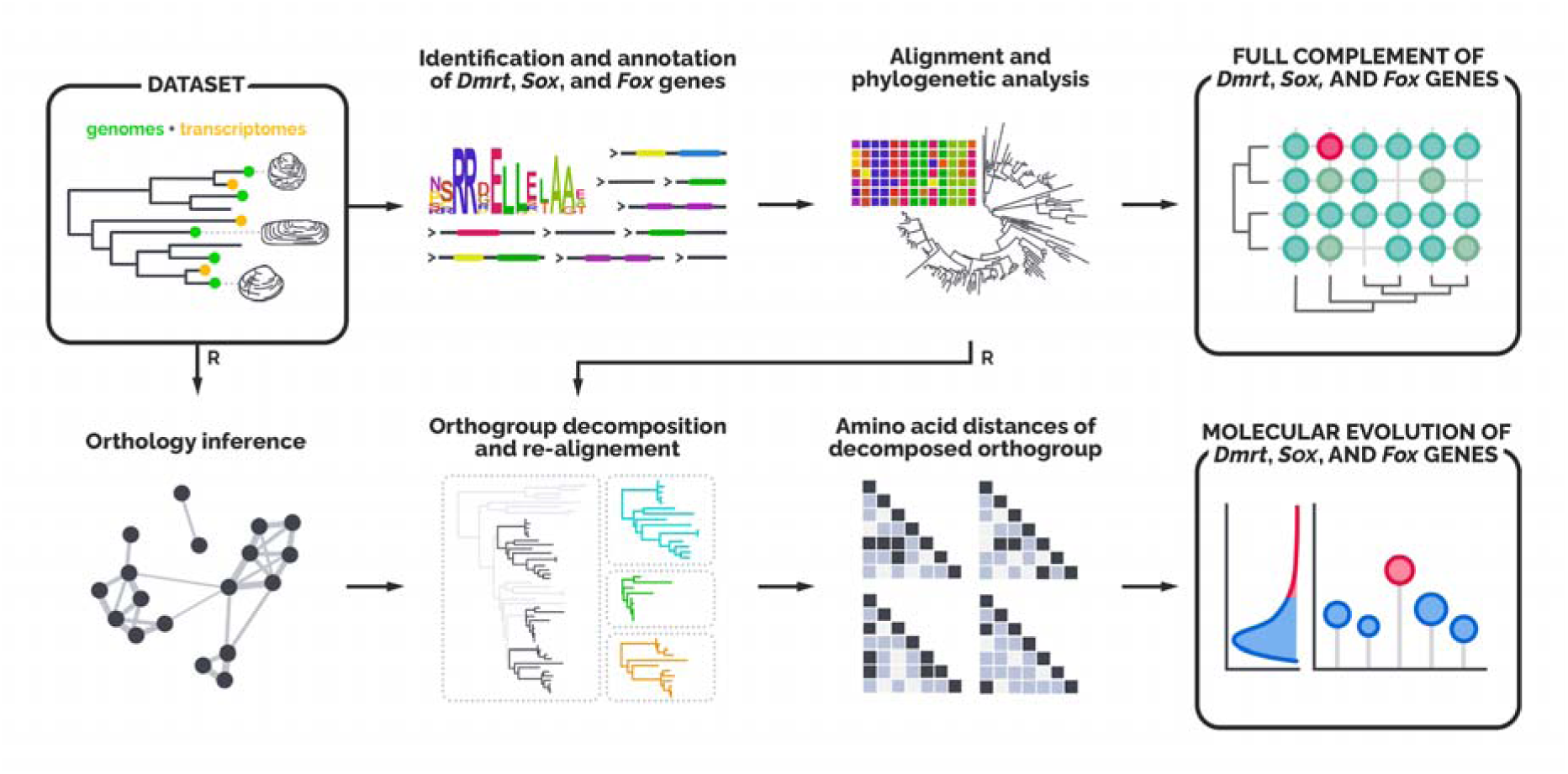
Workflow for the bivalve dataset. Starting from a set of both genomes and transcriptomes covering a great portion of bivalve taxonomic diversity, we first characterized the entire complement of Dmrt, Sox, and Fox genes (upper row). We used sequence annotation and phylogenetic tools to obtain reliable sequences and filter out any putative mis-assembled or mis-annotated sequence. Afterwards, we built a reduced set of transcriptomes and genomes (the reduced bivalve dataset, where we minimized the redundancy of congeneric species) from which to draw the molecular evolution patterns of orthologous genes (bottom row). After having obtained gene single-copy orthologous groups, we calculated the amino acid distances within each orthogroup and then we built the distribution of median values. The same pipeline was also utilized for the mammal and the fruit fly datasets, with just two minor differences: the starting dataset was composed of only genomes, and the reduction step (R) was not necessary.

mtZOA, rtREV, VT, WAG) and the corresponding pairwise amino acid distances were computed with the function ‘dist.ml’ from the ‘phangorn’ R package (**Schliep 2011**). We decided to use the pairwise amino acid distance instead of the tip-to-tip phylogenetic distance (which accounts for a more comprehensive evolutionary signal) to save computational time. However, to check whether the two metrics were comparable to each other, we randomly selected 200 SCOs and computed the maximum likelihood (ML) trees using IQTREE2, with ModelSelection restricted as before. Then, the tip-to-tip pairwise distances were obtained with the R package ‘adephylo’ (**Jombart & Dray, 2010**). The same pipeline to obtain SCOs and AASD was also utilized on the DSFG families. The distribution of AASD was then built after the median values of pairwise distances of each SCO, and genes were categorised accordingly into three groups: Group 1, consisting of genes from the 1% upper quantile of the distribution; Group 2, consisting of genes between the 1% and 5% upper quantiles; and Group 3, consisting of all the remaining genes. Group 1 and Group 2 genes will be referred to as ‘highly divergent genes’.

### Mammals and *Drosophila* spp. as test datasets

To validate our approach for the study of bivalve SRG molecular evolution, we run the same analysis on two additional datasets, consisting of reference genomes of mammals and *Drosophila* species (**Supp. Tab. S4–S5**, respectively), whose sex-determining mechanisms are well studied and characterised. As a matter of fact, despite it is well known that SDGs tend to evolve faster than genes not involved in SD, the hypothesis has never been tested extensively across the entire phylogenetic diversity of a group: molecular evolution of SDGs and SRGs has mainly been tested on single species or inside the boundaries of taxonomic genera (**Stothard et al., 2003**; **Haerty et al., 2007**; **Mank et al., 2007**; **Mullon et al., 2012**; **Papa et al., 2017**; **Ghiselli e al., 2018**). For both mammals and fruit flies, annotated genomes were downloaded from NCBI using the command-line tool ‘datasets’, then processed using the same pipeline and scripts as before (**Fig. 1**).

### GO term enrichment

After having obtained the distributions of AASD in the three datasets (Bivalvia, Mammalia, and *Drosophila*) we performed a gene ontology (GO) enrichment analysis of highly divergent genes. To do so, we firstly selected one gene per SCO, giving priority to few chosen species: (i) for bivalves, we selected genes from *Pecten maximus*, or alternatively from *Crassostrea gigas*, *Hyriopsis bialata*, *Tridacna squamosa*, and *Solen grandis*; (ii) for mammals, we selected genes from *H. sapiens*, or alternatively from *Bubalus bubalis*, *Panthera tigris*, *Camelus dromedarius*, and *Monodelphis domestica*; (iii) for fruit flies, we selected genes from *D. melanogaster*, or alternatively from *Drosophila hydei*, *Drosophila pseudoobscura*, and *Drosophila suzukii*. Afterwards, we annotated the obtained datasets with the corresponding GO terms using the OMA browser (accessed 18/09/2024; **Altenhoff et al., 2024**). The GO-term enrichment was then performed with the R package ‘topGO’ using the Fisher’s exact test (**Alexa et al., 2024**).

## RESULTS

### Assembly of the bivalve, mammal, and fruit fly datasets

The comprehensive bivalve dataset on which we retrieved and analysed DSFGs consists of 29 bivalve genomes, 14 bivalve transcriptomes, and 7 outgroup genomes (5 gastropods and 2 *Octopus* spp.; **Supp. Tab. S1**). BUSCO statistics for complete single-copy genes spanned from the 64.9% in *Modiolus modiolus* to the 99.4% of *Perna viridis*, with a median value of 94.7%. We were able to get at least one representative species for 11 different bivalve orders, covering a good proportion of the phylogenetic diversity of the clades Pteriomorphia, Palaeoheterodonta, and Imparidentia, and thus building the most extensive genomic and transcriptomic dataset for bivalve comparative analyses so far. Unfortunately, no genomes or transcriptomes for Protobranchia, Archiheterodonta, and Anomalodesmata were available at the time of the project, thus we were not able to include any of those clades in our analysis. The mammal dataset consists of 32 species and 1 outgroup (*Gallus gallus*, Aves; **Supp. Tab. S4**), and covers 12 major orders, while the fruit fly dataset consists of 17 species and 1 outgroup (*Anopheles gambiae*, Culicidae; **Supp. Tab. S5**), and covers 2 *Drosophila* subgenera (i.e., *Drosophila* and *Sophophora*). BUSCO statistics for complete single-copy genes were generally higher than those of bivalves, with a median of 98.3% for mammals and of 99.8% for fruit flies (**Supp. Tab. S4–S5**).

### Dmrt, Sox, and Fox complements in bivalves

Our pipeline managed to successfully identify and annotate DSFGs in bivalves, as proved by the same analysis in mammals and fruit flies (see following sections). We retrieved four main orthology groups of Dmrt genes in bivalves (**Fig. 2**; **Supp. Fig. S1**; **Supp. Tab. S6**), three corresponding to the groups present in the Bilateria common ancestor (*Dmrt-2*, *Dmrt-3*, and *Dmrt-4/5*; **Mawaribuchi et al., 2019**), and one additional group with no clear ortholog among reference genes, and thus putatively specific to molluscs (named *Dmrt-1L*, as per **Li et al., 2018b**; **Evensen et al., 2022**). The *Dmrt-4/5* subgroup shows a group-specific expansion in Palaeoheterodonta and Heterodonta, while *Dmrt-1L* is completely absent from Heterodonta. The degree of missing data for Dmrt genes in bivalves is ∼35% (**Supp. Tab. S7**). The ubiquitin-binding CUE-like DMA domain has been annotated in most of the *Dmrt-3* and *Dmrt-4/5* genes, while an additional DM domain has been annotated in *Dmrt-1L* genes in Mytilida and the gastropod *Pomacea canaliculata* (**Supp. Tab. S6**). Additionally, we retrieved six main orthology groups of Sox genes, none of which is restricted to molluscs or bivalves (**Fig. 2**; **Supp. Fig. S2**; **Supp. Tab. S6**). Five Sox groups (*Sox-B1/2*, *Sox-C*, *Sox-D*, *Sox-E*, and *Sox-F*) are those traditionally considered to be present in the Bilateria common ancestor (**Phochanukul & Russell, 2010**), while one has been identified outside mammals only recently (*Sox-H*, or *Sox-30*; **Han et al., 2010**). *Sox-B2* and *Sox-B1* have been grouped in the same clade, as in our phylogenetic reconstruction *Sox-B2* is paraphyletic with respect to *Sox-B1* (**Supp. Fig. S2**), despite being traditionally recognised as a separate paralogy group in humans, fruit flies, and roundworms. The degree of missing data for Sox genes in bivalves is ∼8% (**Supp. Tab. S7**). The Sox N-terminal signature domain was annotated for *Sox-E* genes (**Supp. Tab. S6**). Concerning Fox genes, we retrieved 27 main orthology groups (**Fig. 2**; **Supp. Fig. S3**; **Supp. Tab. S6**), two of which are specific to molluscs (*Fox-OG13/NA*, *Fox-OG16/NA*). Additionally, other potential mollusc-specific Fox groups have been identified, but these have been excluded from the final orthology analysis as they are present in less than half of bivalve species (**Supp. Tab. S6**). The two major Fox gene subgroups, Group I (monophyletic, specific to Metazoa; includes *Fox-A*, *Fox-B*, *Fox-C*, *Fox-D*, *Fox-E*, *Fox-F*, *Fox-G*, *Fox-H*, *Fox-L1*, *Fox-L2*, *Fox-Q2*) and Group II (paraphyletic, specific to Opisthokonta; includes *Fox-O*, *Fox-P*, *Fox-J2*, *Fox-J1*, *Fox-K*, *Fox-N2/3*, *Fox-N1/4*; **Larroux et al., 2008**), have been recovered, including the four Fox genes that were present in the Bilateria common ancestor (*Fox-C*, *Fox-F*, *Fox-L1*, and *Fox-Q1*; **Shimeld et al., 2010**). Two putative lineage-specific expansions have been recovered for *Fox-OG28/NA*, one regarding *Mytilus* spp. and one regarding the two Myida species (**Fig. 2**; **Supp. Fig. S3**). The degree of missing data for Fox genes in bivalves is ∼22% (**Supp. Tab. S7**). The forkhead-associated (FHA) domain was annotated for *Fox-K* genes, the *Fox-P* coiled-coil signature domain was annotated for *Fox-P* genes, while both the forkhead N- and C-terminal signature domains were annotated for *Fox-A* genes (**Supp. Tab. S6**).

**Fig. 2.**
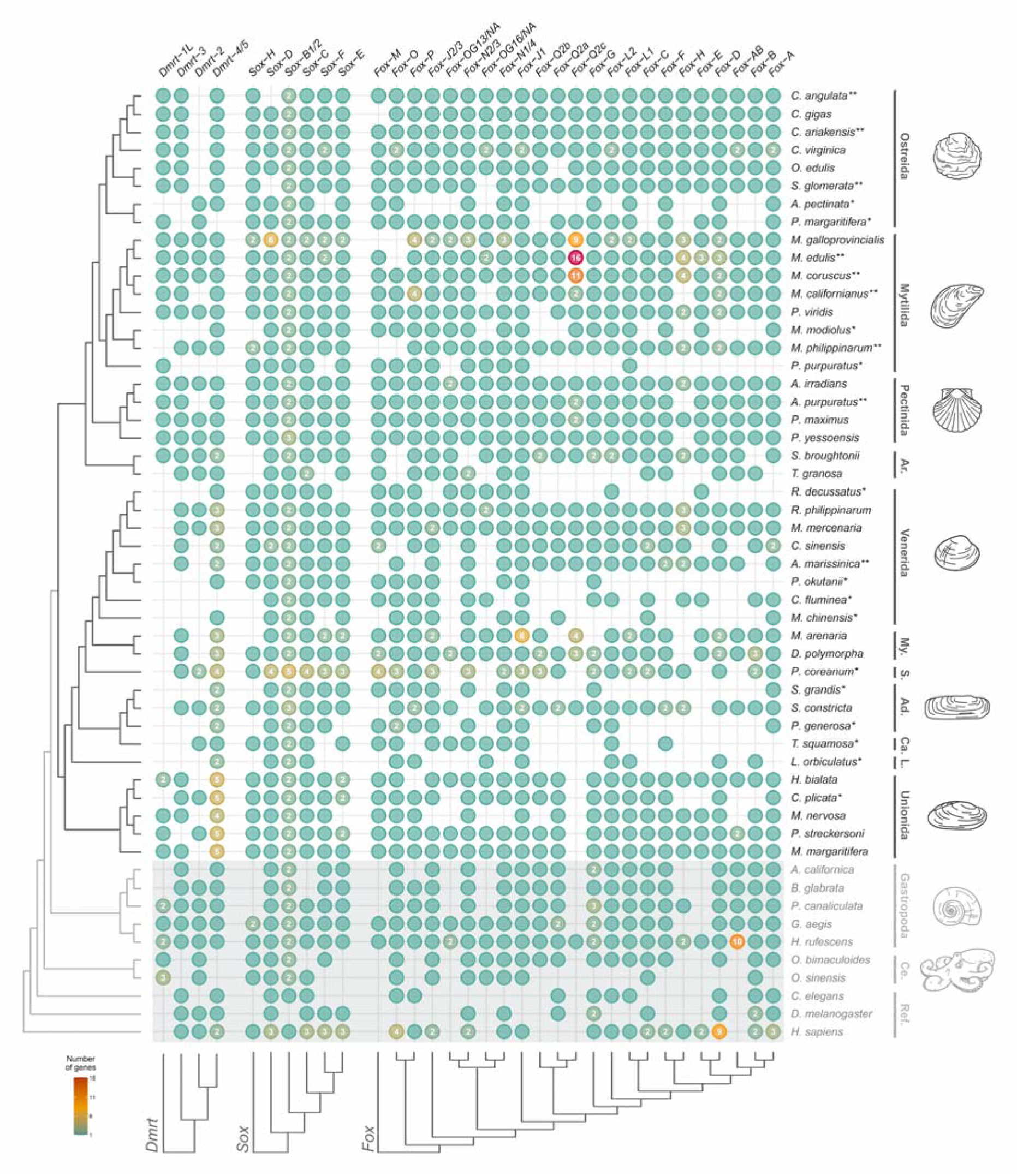
Dmrt, Sox, and Fox gene (DSFG) complement in bivalves and their outgroups. The presence of genes in various species are indicated by filled circles. Numbers inside each circle specify genes with 2 or more copies. The shaded area highlights non-bivalve species, belonging either to other molluscs or to the references. The phylogenetic tree of analyzed species, as inferred from literature, is shown on the left, while major taxonomic groups are reported on the right. Species represented by transcriptomic data are marked with an asterisk (‘*’), and species not present in the reduced bivalve dataset are marked with two asterisks (‘**’; see main text and Fig. 1); note that the two categories do nor overlap. DSFG trees are shown on the bottom (full trees can be found in **Supp. Fig. S1–S3**). Full species names, along with all assembly and taxonomic information, can be found in **Supp. Tab. S1**. Ad.: Adapedonta; Ar.: Arcida; Ca.: Cardiida; Ce.: Cephalopoda; L.: Lucinida; My.: Myida; Ref.: reference genes; S.: Sphaeriida.

### Amino acid sequence divergence of Dmrt, Sox, and Fox genes in bivalves

We produced amino acid alignments of ∼11k SCOs, 32 of which belongs to the DSFG families. Of these, 112 were assigned to Group 1 (1% upper quantile), 447 to Group 2 (5% upper quantile), and 10,603 to Group 3. Most of the DSFGs (29/32) fell in Group 3 (**Fig. 3**), which means they have a median AASD comparable to the vast majority of other genes in bivalves (median level of the genomes). Just *Dmrt-1L*, *Sox-H*, and *Sox-F* showed higher divergences, and have been accordingly placed in Group 2. Overall, pairwise AASD proved to be a good approximation of the tip-to-tip distances calculated on the corresponding ML phylogenetic trees (*R* = 0.84, *p* < 2.2E−16, considering 200 randomly-selected trees; **Fig. 3C**), and it also shows no influence from the alignment length (*R* = 0.11) or the number of represented species (*R* = −0.23; **Fig. 3D–E**). Genes from Group 1 and Group 2 are strongly involved in cellular regulatory processes (such as those related to the metabolism of nucleic acids, proteins, and other macromolecules), but also in development and response to external stimuli, as shown by the GO-term enrichment analysis (**Tab. 1**; **Supp. Tab. S8**).

**Fig. 3.**
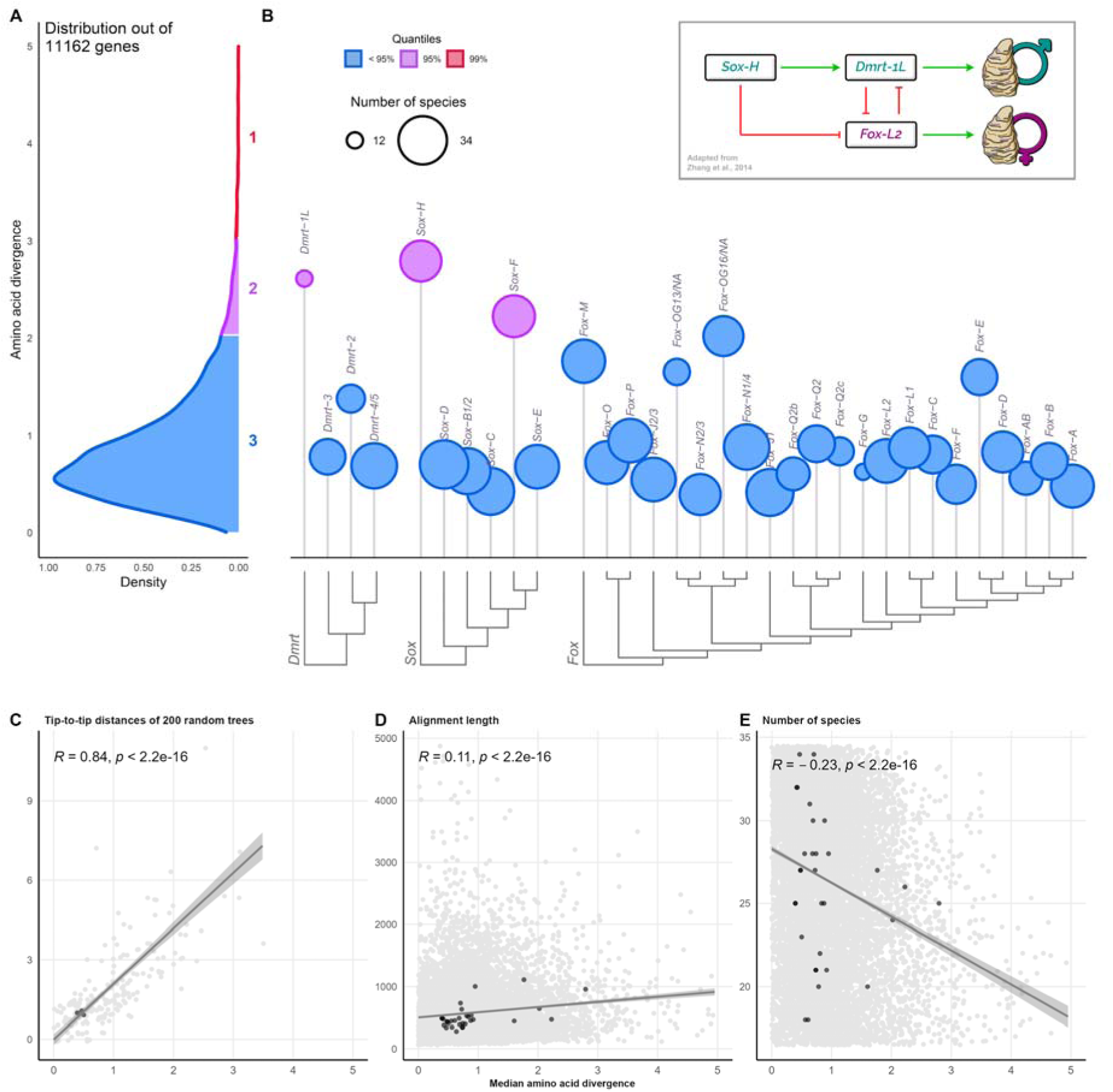
Distribution of amino acid sequence divergence (AASD) of single-copy orthogroups in bivalves (A), including Dmrt, Sox, and Fox genes (DSFGs; B), and their correlations with tip-to-tip distances (C), alignment lengths (D), and number of species (E). The distribution of AASD has been computed on the median values of pairwise distances of over 11k SCOs from the reduced bivalve dataset (see main text and Fig. 1). Genes have been divided according to their median AASD value into three different groups, which are indicated by different colours and increasing numbers (Groups 1, 2, and 3). Genes from Group 1 and Group 2 are collectively named ‘highly divergent genes’. Circle heights of DSFGs show the median value of their AASD, while the size indicates the number of represented species. DSFG trees are shown on the bottom (full trees can be found in **Supp. Fig. S1–S3**). Darker points in C-E indicate Dmrt, Sox, and Fox gene SCOs. The correlation between the amino acid distance and the tip-to-tip distance has been computed on 200 randomly selected orthogroups. **Insets:** scheme of the sex-determination working model in *Crassostrea gigas* as hypothesised by **Zhang et al. (2014)**, showing the main genes involved. Green arrows indicate transcription activations, red arrows indicate transcription suppressions.

**Tab. 1.** Top enriched GO terms for highly divergent genes of bivalves, mammals, and *Drosophila*. The extended version of the table, which includes also the expected number of annotated genes per GO term and all the other enriched GO terms, can be accessed in **Supp. Tab. S8**.

| Dataset | GO.ID | Term | Annotated genes | Significant genes | Corrected p-value |
| --- | --- | --- | --- | --- | --- |
| **Bivalvia** | GO:0060255 | regulation of macromolecule metabolic process | 737 | 59 | 0.04525 |
|  | GO:0080090 | regulation of primary metabolic process | 673 | 53 | 0.01818 |
|  | GO:0019219 | regulation of nucleobase-containing compound metabolic process | 541 | 41 | 0.02388 |
|  | GO:0006351 | DNA-templated transcription | 571 | 39 | 0.03767 |
|  | GO:0032774 | RNA biosynthetic process | 579 | 39 | 0.04490 |
|  | GO:0051252 | regulation of RNA metabolic process | 517 | 37 | 0.02719 |
|  | GO:0006355 | regulation of DNA-templated transcription | 490 | 35 | 0.03751 |
|  | GO:2001141 | regulation of RNA biosynthetic process | 491 | 35 | 0.03844 |
|  | GO:0006950 | response to stress | 370 | 33 | 0.01949 |
|  | GO:0032502 | developmental process | 261 | 27 | 0.04445 |
|  | GO:0006468 | protein phosphorylation | 345 | 23 | 0.02483 |
|  | GO:0031325 | positive regulation of cellular metabolic process | 125 | 17 | 0.00801 |
|  | GO:0010604 | positive regulation of macromolecule metabolic process | 151 | 17 | 0.04047 |
|  | GO:0051172 | negative regulation of nitrogen compound metabolic process | 117 | 16 | 0.00814 |
|  | GO:0051173 | positive regulation of nitrogen compound metabolic process | 137 | 15 | 0.02454 |
|  | GO:0006310 | DNA recombination | 66 | 14 | 0.00087 |
|  | GO:0048513 | animal organ development | 83 | 12 | 0.04088 |
|  | GO:0010629 | negative regulation of gene expression | 78 | 11 | 0.00048 |
|  | GO:0023051 | regulation of signaling | 133 | 11 | 0.02872 |
|  | GO:0045934 | negative regulation of nucleobase-containing compound metabolic process | 64 | 11 | 0.03637 |
|  | GO:0009605 | response to external stimulus | 90 | 11 | 0.04544 |
|  | GO:0044419 | biological process involved in interspecies interaction between organisms | 63 | 11 | 0.04761 |
| **Mammalia** | GO:0006955 | immune response | 1297 | 145 | 0.00061 |
|  | GO:0098542 | defense response to other organism | 853 | 112 | 0.02066 |
|  | GO:0045087 | innate immune response | 647 | 82 | 8.5e-10 |
|  | GO:0001817 | regulation of cytokine production | 630 | 51 | 0.04660 |
|  | GO:0042742 | defense response to bacterium | 233 | 45 | 1.7e-07 |
|  | GO:0006954 | inflammatory response | 642 | 45 | 0.01735 |
|  | GO:0019221 | cytokine-mediated signaling pathway | 382 | 44 | 3.9e-07 |
|  | GO:0002250 | adaptive immune response | 342 | 44 | 1.3e-05 |
|  | GO:0001819 | positive regulation of cytokine production | 402 | 41 | 0.02723 |
|  | GO:0002697 | regulation of immune effector process | 308 | 37 | 0.04426 |
|  | GO:0042110 | T cell activation | 432 | 35 | 0.02564 |
|  | GO:0051607 | defense response to virus | 257 | 34 | 1.9e-07 |
|  | GO:0048232 | male gamete generation | 491 | 32 | 0.02255 |
|  | GO:0007283 | spermatogenesis | 478 | 31 | 0.02801 |
|  | GO:0070661 | leukocyte proliferation | 273 | 29 | 0.01285 |
|  | GO:0002449 | lymphocyte mediated immunity | 221 | 29 | 0.04833 |
|  | GO:0070663 | regulation of leukocyte proliferation | 212 | 25 | 0.01870 |
|  | GO:0050727 | regulation of inflammatory response | 300 | 24 | 0.00235 |
|  | GO:0031349 | positive regulation of defense response | 240 | 24 | 0.01239 |
|  | GO:0002768 | immune response-regulating cell surface receptor signaling pathway | 177 | 22 | 0.00336 |
|  | GO:0050829 | defense response to Gram-negative bacterium | 66 | 17 | 1.7e-10 |
|  | GO:0071222 | cellular response to lipopolysaccharide | 164 | 17 | 0.00012 |
|  | GO:0010466 | negative regulation of peptidase activity | 163 | 16 | 0.00036 |
|  | GO:0002429 | immune response-activating cell surface receptor signaling pathway | 164 | 16 | 0.00243 |
|  | GO:1903555 | regulation of tumor necrosis factor superfamily cytokine production | 137 | 16 | 0.01244 |
|  | GO:0071706 | tumor necrosis factor superfamily cytokine production | 137 | 16 | 0.01244 |
|  | GO:0070665 | positive regulation of leukocyte proliferation | 132 | 16 | 0.02765 |
|  | GO:0045089 | positive regulation of innate immune response | 113 | 16 | 0.03224 |
|  | GO:0071356 | cellular response to tumor necrosis factor | 175 | 15 | 0.00219 |
|  | GO:0002695 | negative regulation of leukocyte activation | 148 | 15 | 0.01151 |
|  | GO:0002456 | T cell mediated immunity | 82 | 15 | 0.01605 |
|  | GO:0002705 | positive regulation of leukocyte mediated immunity | 113 | 15 | 0.01837 |
|  | GO:0032680 | regulation of tumor necrosis factor production | 133 | 15 | 0.03262 |
|  | GO:0032640 | tumor necrosis factor production | 133 | 15 | 0.03262 |
|  | GO:0050866 | negative regulation of cell activation | 165 | 15 | 0.04048 |
| Drosophila | GO:0000819 | sister chromatid segregation | 140 | 11 | 0.02927 |
|  | GO:0070192 | chromosome organization involved in meiotic cell cycle | 54 | 9 | 0.00849 |
|  | GO:0007131 | reciprocal meiotic recombination | 37 | 7 | 0.00066 |
|  | GO:0007143 | female meiotic nuclear division | 54 | 6 | 0.02270 |
|  | GO:0035967 | cellular response to topologically incorrect protein | 44 | 5 | 0.03334 |
|  | GO:0035966 | response to topologically incorrect protein | 47 | 5 | 0.04266 |
|  | GO:0007141 | male meiosis I | 13 | 4 | 0.00150 |
|  | GO:0140543 | positive regulation of piRNA transcription | 3 | 3 | 6.9e-05 |
|  | GO:0010526 | retrotransposon silencing | 8 | 3 | 0.00331 |
|  | GO:0007130 | synaptonemal complex assembly | 10 | 3 | 0.00666 |
|  | GO:0030719 | P granule organization | 11 | 3 | 0.00888 |
|  | GO:0071218 | cellular response to misfolded protein | 12 | 3 | 0.01149 |
|  | GO:0051788 | response to misfolded protein | 12 | 3 | 0.01149 |
|  | GO:0007135 | meiosis II | 15 | 3 | 0.02169 |
|  | GO:0034508 | centromere complex assembly | 19 | 3 | 0.04094 |

### Dmrt, Sox, and Fox genes, and amino acid sequence divergence in the test datasets

Most of the already-recognised DSFG orthology groups in mammals and fruit flies have been identified. In mammals, we retrieved 7 Dmrt orthology groups with ∼3.1% of missing data, 20 Sox orthology groups with ∼8.1% of missing data, and 42 Fox orthology groups with ∼4.6% of missing data (**Supp. Fig. S4A, S5–S7**; **Supp. Tab. S9**). Of these, just *Sox-5* was not included in the subsequent AASD analysis, as it did not meet the 50%-species occupancy threshold. OrthoFinder analysed ∼650M genes, and the number of SCOs used in the AASD analysis is >16k (**Fig. 4A**). From the distribution of median AASD, 163 genes were assigned to Group 1, 649 to Group 2, and 15,355 to Group 3. Most of the DSFGs (66/68) fell in Group 3 (**Fig. 4B**), while *Sry* and *Fox-D4* showed higher divergences, and have been accordingly placed in Group 1 and 2, respectively. Genes from Group 1 and Group 2 show a strong enrichment in immune-related functions (such as innate and adaptive immune response, defence response to bacteria and viruses, lymphocyte metabolism, etc.), but also in reproductive processes (such as spermatogenesis; **Tab. 1**; **Supp. Tab. S8**). Concerning *Drosophila*, we retrieved 4 Dmrt orthology groups with ∼1.7% of missing data, 7 *Sox* orthology groups with ∼3.9% of missing data, and 17 Fox genes with ∼8.3% of missing data (**Supp. Fig. S4B, S9–S10**; **Supp. Tab. S10**). OrthoFinder analysed ∼240M, and the distribution of median AASD was built after >12k SCOs (**Fig. 4C**). 126 genes were assigned to Group 1, 501 to Group 2, and 11,880 to Group 3. All the DSFGs have been used in the AASD analysis, but none of them have been placed in Group 1 or 2, that is, all the DSFGs in *Drosophila* have an AASD comparable to the median level of the genome (**Fig. 4D**). Genes of Group 1 and Group 2 show a GO-term enrichment in meiotic processes, such as chromosome/chromatid organisation, and retrotransposon silencing (**Tab. 1**; **Supp. Tab. S8**).

**Fig. 4.**
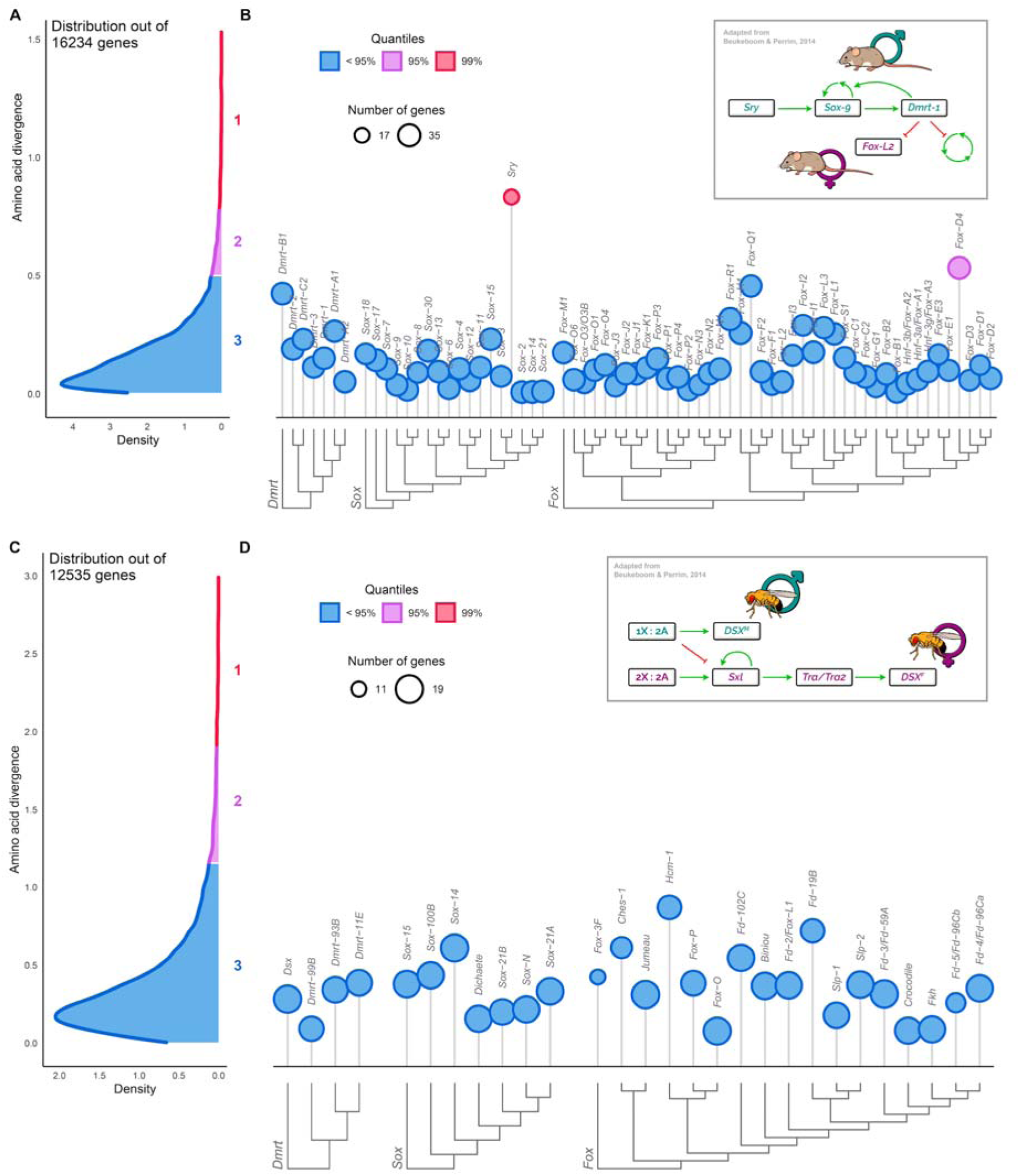
Distribution of amino acid divergence (AASD) of single-copy orthogroups in Mammalia (A) and *Drosophila* (C), including Dmrt, Sox, and Fox genes (DSFGs; B-D). The distributions of AASD in mammals and fruit flies have been computed on the median values of pairwise distances of over 16k and 12k SCOs, respectively. Genes have been divided according to their median AASD value into three different groups, which are indicated by different colors and increasing numbers (Groups 1, 2, and 3). Genes from Group 1 and Group 2 are collectively named ‘highly divergent genes’. Circle heights of DSFGs show the median value of their AASD, while the size indicates the number of represented species. DSFG trees are shown on the bottom (full trees can be found in **Supp. Fig. S5–S7** for mammals and in **Supp. Fig. S8–S10** for fruit flies). In *Drosophila*, *Sxl* and *tra*, both involved in the SD pathway (inset) do not belong to the group of highly divergent genes (mean amino acid divergence of 0.0923793827106834 and 0.978190195613595, respectively; that is, within the boundaries of Group 3). **Insets:** scheme of the sex-determination molecular pathways in *Mus musculus* and in *Drosophila melanogaster*, showing the main genes involved (adapted from **Beukeboom & Perrim, 2014**). Green arrows indicate transcription activations, red arrows indicate transcription suppressions. X: sex chromosomes; A: autosomal chromosomes; *DSX^M/F^*: *DSX* splicing variants present in males or females, respectively.

## DISCUSSION

### A new manually-curated and phylogenetic-based reference dataset of Dmrt, Sox, and Fox genes in bivalves

The annotation and characterisation process of a gene family may harbour many overlooked challenges in a certain clade of organisms (**Vizueta et al., 2020**). For example, the presence of highly conserved catalytic domains may hamper the correct identification of the components of a gene family because of insufficient phylogenetic signal, as it is the case for Hox and ParaHox genes and their homeobox motif (**Baldwin-Brown et al., 2018**; **Nicolini et al., 2023b**). Conversely, the components of dynamic gene families characterised by abrupt and sequential duplication events may be difficult to sort into separate groups. As a matter of fact, varying levels of sequence heterogeneity and gene copy numbers makes the inference of orthologous groups hard, as for certain clans of the P450 family (**Dermauw et al., 2020**). Regardless of the causes, having a solid and wide phylogenetic context, in which to study gene duplications/losses and orthology relationships, is crucial to overcome these difficulties. In the same way, manual curation and visual inspection of multiple sequence alignments, phylogenetic trees, and gene structures (in terms of domain composition, start and stop codons, and other feature representations) is helpful, despite being time-demanding and possibly less reproducible. In this study, we characterised the full complement of DSFGs in the vast class of bivalves, by leveraging sequence domain annotation, phylogenetics, and manual curation of the dataset. Our aim was to obtain the most reliable gene complements as possible, combined with a vast taxonomic dataset, a solid phylogenetic inference, an openly available dataset of gene sequences, and a reproducible pipeline for the annotation of gene identity. By doing so, we want to provide a reliable resource for future studies of DSFGs, either focused on bivalves or on Metazoa in general.

Concerning the Dmrt gene family, we identified orthologs of the vertebrate *Dmrt-2*, *Dmrt-3*, and *Dmrt-4/5* (*A1/A2;* **Fig. 2**; **Supp. Fig. S1**; **Supp. Tab. S6**), which are also expected to have been present in the Bilateria common ancestor (**Mawaribuchi et al., 2019**). **Wang et al. (2023)** found that *Dmrt-4/5* is duplicated in *Mercenaria mercenaria* and *Cyclina sinensis* (Venerida), and in *Dreissena polymorpha* (Myida), and we confirm this result by tracing back the duplication event to the split between Palaeoheterdonta (here represented by Unionida) and Heterodonta (here represented by Venerida, Myida, Sphaeriida, Adapedonta, Cardiida, and Lucinida; **Fig. 2**). Furthermore, we confirm *Dmrt-1L* to be present in many bivalve species (mainly belonging to the Ostreida, Pectinida, Mytilida, and Unionida orders; **Fig. 2**), as well as in gastropods and *Octopus*. Though, our phylogenetic analysis did not retrieve any unambiguous orthology relationship among *Dmrt-1L* and either vertebrate *Dmrt-1* or *Drosophila dsx* genes, as instead it was proposed in previous works (**Li et al., 2018b**; **Evensen et al., 2022**). As a matter of fact, the amino acid sequence of the *Dmrt-1L* DM domain does not recall that of any other Dmrt gene. Furthermore, it must be considered that various phylogenetic analyses have recovered both *Dmrt-1* and *dsx* genes to be restricted to vertebrates and arthropods, respectively (**Wexler et al., 2014**; **Mawaribuchi et al., 2019**; **Panara et al., 2019**), that is, they do not have any direct ortholog outside their relative clades. Thus, if *Dmrt-1L*, *dsx*, and *Dmrt-1* were true orthologs, their origin would need to be placed at least in the Bilateria common ancestor, which seems however to be not the case. All considered, we thus confirm that *Dmrt-1L* is not orthologous to *Dmrt-1* and *dsx* and is rather a mollusc-specific gene (**Evensen et al., 2022**). The monophyly of the *Dmrt-1L* group is not supported by the phylogenetic tree inferred with Dmrt genes from molluscs and the reference species (**Supp. Fig. S1**); though, it is recovered when analysing just genes from mollusc species (**Supp Fig. S11**). To this regard, we speculate that in our analysis, the difficulty in obtaining the monophyly of *Dmrt-1L* genes may have arisen primarily because of the many *C. elegans*-restricted genes (**Supp. Tab. S3**), which are placed among bivalve genes (**Supp. Fig. S1**), but also because of the high AASD of *Dmrt-1L* genes (see **High amino acid sequence divergence identifies putative sex-determining genes**), which hampers a straight-forward phylogenetic reconstruction. Furthermore, our broad-context analysis allowed us to identify some cases of incorrect gene identification in bivalves, which have arisen because of erroneous or ambiguous annotations in previous works, as a result of limited datasets or analyses. For example, (i) the scallop-specific cluster of Dmrt genes retrieved by **Wang et al. (2023)** rather belongs to the *Dmrt-1L* group, and (ii) the classification of Dmrt genes in *Crassostrea* species provided by **Zeng et al. (2024)** needs to be revised following as follows: *Dmrt-1* genes are *Dmrt-4/5*; *Dmrt-2* genes are *Dmrt-3*; *Dmrt-3* genes are *Dmrt-1L*; hence, *Crassostrea* species do not have *Dmrt-2* genes.

For what concerns the Sox gene family, bivalves (or molluscs) do not show any major clade-restricted gene, as only the five Bilateria-specific Sox groups (*Sox-B1/2*, *Sox-C*, *Sox-D*, *Sox-E*, and *Sox-F*) and *Sox-H* have been identified (**Fig. 2**; **Supp. Fig. S2**; **Supp. Tab. S6**), in accordance with previous findings (**Yu et al., 2017b**; **Evensen et al., 2022**; **Wang & Nie, 2024**). *Sox-B1/2* is clearly made up of two subgroups (i.e., *Sox-B1* and *Sox-B2*), as expected, but their respective identity could not be unambiguously established, as *Sox-B1/2* genes of reference species do not form separate clusters (**Supp. Fig. S2**), even when inferring the phylogenetic tree only of components of the *Sox-B1/2* group from molluscs and reference species (**Supp. Fig. S12**).

Compared to Dmrt and Sox genes, the Fox gene family appears as the most dynamic in terms of gene presence/absence, as already shown by other works (**Wu et al., 2020**; **Schomburg et al., 2022**; **Seudre et al., 2022**). Our phylogenetic analysis successfully recovered Group I and Group II of Fox genes (**Larroux et al., 2008**), which include the four Fox genes that were present in the Bilateria common ancestor (*Fox-C*, *Fox-F*, *Fox-L1*, and *Fox-Q1*; **Fig. 2**; **Supp. Fig. S3**; **Supp. Tab. S6**; **Shimeld et al., 2010**). To our knowledge, this is the first broad-taxonomic identification and classification of Fox genes in bivalves, as up to now they have been systematically characterised only in *C. gigas* (**Yang et al., 2014**), *Patinopecten yessoensis* (**Wu et al., 2020**), and *Ruditapes philippinarum* (**Liu et al., 2024**). Firstly, our analysis confirms the absence in molluscs of *Fox-I*, *Fox-Q1*, *Fox-R*, *Fox-S* (**Supp. Fig. S3**), which are in fact thought to have emerged with the diversification of deuterostomes or vertebrates (**Yang et al., 2014**; **Wu et al., 2020**; **Schomburg et al., 2022**; **Seudre et al., 2022**). Furthermore, we have found many Fox groups that appeared as mollusc-specific and/or still-unnamed at a first analysis. However, a more in-depth investigation revealed a different scenario. *Fox-OG2/NA* appears close to the human *Fox-M* gene in the phylogenetic tree, but they do not form a monophyletic group (**Supp. Fig. S3**). However, by comparing *Fox-OG2/NA* sequences and phylogenetic tree with those analysed by **Yang et al. (2014)**, **Wu et al., (2020)**, **Schomburg et al. (2022)**, **Seudre et al. (2022)**, it appears clear that this group of Fox genes is indeed *Fox-M*. However, our analysis has failed to retrieve a monophyletic relationship among bivalve and human *Fox-M* genes, even when inferring a tree with just *Fox-J2*, *Fox-M*, *Fox-O*, and *Fox-P* complements (**Supp. Fig. S13**), which belong to the same Fox group. Regarding the *Fox-OG39/NA* group, it does not have any homolog in reference species (**Supp. Fig. S3**) but is found to belong to the *Fox-AB* group by sequence comparison with previous works (**Yang et al., 2014**; **Wu et al., 2020**; **Seudre et al., 2022**). *Fox-AB* was formerly described only in the sea urchin *Strongylocentrotus purpuratus* and the lancelet *Branchiostoma floridae* (**Tu et al., 2006**; **Yu et al., 2008**), but was later identified also in several Spiralia lineages, including molluscs (e.g., **Yang et al., 2014**; **Wu et al., 2020**; **Seudre et al., 2022**). A similar situation concerns *Fox-OG15/NA* and *Fox-OG28/NA*, which again could not be named based on orthology relationships with the reference species genes (**Supp. Fig. S3**), but actually represent two lineage-specific expansions of the *Fox-Q2* group (named *Fox-Q2b* and *Fox-Q2c*), as already appointed in previous studies (**Yang et al., 2014**; **Wu et al., 2020**). This observation fits within the wider context of the *Fox-Q2* group expansion in Bilateria and, particularly, in Spiralia, that led to remarkable differences in their gene copy numbers across various clades (**Seudre et al., 2022**). Two additional Fox genes have been previously identified in bivalves and were named *Fox-Y* and *Fox-Z* (**Yang et al., 2014**; **Wu et al., 2020**). In our analysis, these Fox genes were identified as *Fox-OG13/NA* and *Fox-OG16/NA*, after sequence comparison with Fox genes from *C. gigas* and *P. yessoensis*. On one hand, *Fox-Y* was firstly identified in *S. purpuratus* (**Tu et al., 2006**) and only recently in a few bivalve species (**Yang et al., 2014**; **Wu et al., 2020**). However, when analysing bivalve and *S. purpuratus* Fox genes, we failed in retrieving such a clear orthology relationship, as *S. purpuratus Fox-Y* does not fall within the phylogenetic range of bivalve *Fox-OG13/NA*, which contains the supposed *Fox-Y* orthologs (**Supp. Fig. S14**). Also, the forkhead domains of *Fox-OG13/NA* genes were annotated as ‘forkhead domain P’ (**Supp. Tab. S6**). On the other hand, *Fox-Z* was firstly identified in bivalves and in several other protostomes, thanks to a phylogenetic work including the brachiopod *Lingula unguis*, the annelid *Capitella teleta*, the scorpion *Centruroides sculpturatus*, and the centipede *Strigamia maritima* (**Wu et al., 2020**).

However, later works have not recovered this Fox gene, even when analysing annelids (**Seudre et al., 2022**) and panarthropods (**Schomburg et al., 2022**) in a more focused effort. In this case, the forkhead domains were annotated as either a generic ‘forkhead domain’ or a ‘forkhead domain Q2’ (**Supp. Tab. S6**). All considered, we argue that bivalves possess two additional Fox groups (here *Fox-OG13/NA* and *Fox-OG16/NA*; **Fig. 2**; **Supp. Fig. S3**; **Supp. Tab. S6**) which are shared with other mollusc species, as revealed also by other authors. However, given the discordant results of the phylogenetic hypothesis and domain annotation, we think that a more thorough investigation on their orthology relationships with Fox genes from other Metazoa is needed, and thus we chose to not employ their former names *Fox-Y* and *Fox-Z*.

Besides the DSFG groups discussed so far, it must be also considered that many orphan genes have been identified (**Supp. Fig. S1–S3**; **Supp. Tab. S6**). For example, **Wu et al. (2020)** identified a duplication event of *Fox-H* genes in *C. gigas*, which has been recovered also in our analysis for the entire Ostreida clade (*Fox-OG36/NA*; **Supp. Fig. S3**). Similarly, a gene orthology group putatively specific to Pteriomorphia has been identified among Sox genes (*Sox-OG1/NA*). Of course, these genes deserve as much attention as their widely distributed paralogs, as they may constitute true group-specific expansions and may play fundamental roles in some biological processes. However, they have not been discussed here or included in **Fig. 2** for clarity purposes, but their sequences are available in on GitHub (see the **Data Availability Statement**).

Overall, our analysis clearly shows the importance of adopting a wide-angle approach when characterising the members of a gene family, especially for large ones such as the Fox genes (**Schomburg et al., 2022**). As a matter of fact, the presence of duplication events and orphan genes needs to be addressed with a broad taxonomic dataset, in order to account for possible mis-annotations, gene phylogenetic misplacements, and sequence heterogeneity. Additionally, many reference species need to be included for the gene identification process, to consider distantly related genes and obtain a solid annotation. Our gene annotation pipeline also resulted to be very solid, even with non-model organisms and sub-optimal genomic and transcriptomic resources as those of bivalves. As a matter of fact, by running the same pipeline on two additional datasets composed of mammal and fruit fly genomes, we were able to obtain high-quality orthology groups in accordance with previous knowledge on the clades (**Supp. Fig. S4-S10**; **Supp. Tab. S9–S10**), with little or no need of manual curation. Furthermore, the present work represents the first broad analysis of DSFGs in both mammals and fruit flies, as so far attention has been mainly dedicated to single well-studied organisms or little clades (e.g., **Jackson et al., 2010**).

### High amino acid sequence divergence identifies putative sex-determining genes

Sex-biased genes tend to evolve more rapidly than unbiased genes at the level of their protein sequences. Accelerated rates have been observed in both male-biased genes (reviewed in **Parsch and Ellegren 2013**; **Grath and Parsch 2016**) and female-biased genes (e.g., **Papa et al. 2017**; **Ghiselli et al. 2018**), but also in SRGs and SDGs (**O’Neil & Belote, 1992**; **Whitfield et al., 1993**; **de Bono & Hodgkin, 1996**). For example, it has been shown that *Dm-W*, *Dmy*, and *Sry* (which are SDGs in the African clawed frog *X. laevis*, in the medaka fish *O. latipes*, and in eutherians, respectively) all have higher substitution rates than their paralogues (*Dmrt-1* for *Dm-W* and *Dmy*, *Sox-3* for *Sry*), particularly when considering their DNA-binding domains (**Mawaribuchi et al. 2012**). Similarly, both a burst of positive selection and a relaxation of purifying selection has been detected in *Drosophila Sxl* in correspondence with its recruitment at the top of the sex-determing cascade. The same signs of relaxed purifying selection have been found in the downstream targets of *Sxl*, that is, *transformer* (*tra*) and *dsx*, despite no evidence of positive selection has been detected (**Mullon et al., 2012**).

Considering these shared features of SRGs and SDGs, we decided to look for signs of accelerated sequence evolution in DSFGs of bivalves to evaluate if any of them could be *a priori* associated with SD by employing the tools of molecular evolution. However, we analysed patterns of sequence evolution not only among putative SRGs and their close paralogs, but also considering the genomic context in which these genes evolve. Our aim was to check whether SRGs show higher rates of sequence evolution when compared to other genes not involved in SD and not belonging to the same gene family. To do so, we obtained the AASD median values of more than 11k SCOs from bivalve genomes (**Fig. 3A**), in order to build a statistical distribution to be used as a reference: if SRGs/SDGs (in this case, DSFGs) truly evolve faster than other genes, we may expect them to fall within the 5% (or even 1%) upper quantile of the distribution (**Fig. 3B**), i.e., within highly divergent genes (Group 1 and Group 2 genes of the distribution). We chose to use the AASD as a metric of sequence evolution (instead of the tip-to-tip distances of phylogenetic trees, which account for more comprehensive evolutionary models) to save computational time. As a matter of fact, the AASD median values proved to be a good approximation of the tip-to-tip median distances in 200 randomly selected genes (**Fig. 3C**; R = 0.84, p < 2.2E−6).

Among DSFGs, three fell within the 5% upper quantile, namely *Dmrt-1L*, *Sox-H*, and *Sox-F*. Interestingly, on the basis of DGE analyses, *Dmrt-1L* and *Sox-H* have been already proposed to be involved in the male SD pathway of *C. gigas* (inset in **Fig. 3B**; **Zhang et al., 2014**), while *Sox-H* to be also a candidate gene involved in the development of male germ cells in the Manila clam *Ruditapes philippinarum* (**Ghiselli et al., 2012**). Specifically, *Sox-H* would play a major role in *C. gigas* SD, by interacting with *Dmrt-1L* and determining the onset of the male phenotype development; at the same time, both *Sox-H* and *Dmrt-1L* would inhibit *Fox-L2*, which instead is necessary to start the female phenotype development. *Dmrt-1L* and *Sox-H* have been appointed several other times to be involved in male-gonad development and differentiation, through DGE (e.g., **Teaniniuraitemoana et al., 2014**; **Capt et al, 2018**; **Afonso et al., 2019**), ISH (e.g., **Naimi et al., 2009**; **Li et al., 2018**; **Liang et al., 2019**; **Yue et al., 2021**), and RNA interference (**Liang et al., 2019**; **Sun et al., 2022**). Therefore, the high AASD of *Dmrt-1L* and *Sox-H* is coherent with previous works, strengthening their role as putative SRGs.

The relationship between high gene AASD and the involvement in SD is particularly strengthened when looking at the patterns of AASD in the test datasets, which corroborates the solidity of our analysis: (i) in the mammal dataset—which represents a strictly genetic SD system with a master and rapidly-evolving SDG—one of the genes from the 5% upper quantile of the distribution is *Sry* (**Fig. 4A–B**), the male sex-determining gene in eutherians (inset in **Fig. 4B**); (ii) in the fruit fly dataset—which represents a chromosomic SD system and with no differences in the rates of sequence evolution among SRGs—none of the DSFG exhibit significantly higher AASD (**Fig. 4C–D**), including the downstream effector *dsx* (inset in **Fig. 4D**). Also, *Sxl* and *tra*, both involved in the SD pathway of *Drosophila* (inset in **Fig. 4D**) do not belong to the group of highly divergent genes, as they have a mean amino acid divergence of about 0.09 and 0.9, respectively (Group 3; **Fig. 4D**). Therefore, it can be argued that both *Dmrt-1L* and *Sox-H* may not only be SRGs, but may participate in bivalve SD as primary SDGs, which is reflected in their high AASD, as it is observed for *Sry* in mammals. As a matter of fact, if they were involved in SD just as intermediate actors of the signalling cascade, we should have not observed a high AASD, as *Drosophila Sxl*, *tra*, and *dsx* seem to suggest. Overall, these patterns of molecular evolution concerning SRGs and SDGs are also supported by the way SD regulatory networks evolve. It has been proposed that the sex-determining cascades tend to arise and be established with a bottom-up mechanism (**Wilkins et al., 1995**; **Mullon et al., 2012**; **Beukeboom & Perrim, 2014**; **Capel, 2017**). This means that the regulatory relationships among genes at the bottom of the cascade are settled prior to the regulatory relationships among genes at the top and, consequently, upstream regulators are progressively recruited to fine-tune diverse SD signals. These evolutionary patterns eventually produce gene regulatory networks in which the divergence of the upstream triggers is higher than that of downstream effectors, in terms of both identity and sequence composition (**Beukeboom & Perrim, 2014**). This mechanism has been proposed for *Drosophila* species (**Mullon et al., 2012**), *C. elegans* (**Stothard & Pilgrim, 2003**), and vertebrates, despite in the latter case it has been questioned several times (reviewed in **Capel, 2017**).

Two main objections can be moved against our approach: (1) the distribution of AASD is not appropriate for this kind of inference, as it does not represent the true gene evolutionary (or substitution) rates (which instead are those usually utilized when dealing with SRGs and SDGs); (2) the three datasets are not comparable one to each other, as they take into consideration very different animal groups, with different taxonomic rankings and different divergence times (thus, the patterns of AASD may be the products of other confounding factors not directly related to SD). Concerning the first objection, we are aware that the AASD does not represent the evolutionary rate itself, but rather its product. However, the two features are tightly linked, as on the long term highly divergent proteins tend to be produced by genes with high evolutionary (or substitution) rates (**Echave et al., 2016**). By performing a GO term enrichment, it emerged that highly divergent genes of the mammal dataset are mainly involved in the immune response and male spermatogenesis (**Tab. 1**; **Supp. Tab. S8**), which are two processes notoriously connected with rapid sequence evolution (i.e., higher evolutionary rates; **Swanson & Vacquier, 2002**; **Murat et al., 2023**; **Vinkler et al., 2023**). Similarly, highly divergent genes from the fruit fly dataset show an enrichment for GO terms associated with meiotic-related functions (such as the formation of the synaptonemal complex by the products of *c(2)M*, *c(3)G*, *corona*, and *corolla* genes; **Tab. 1**; **Supp. Tab. S8**), which again are known to evolve rapidly (**Hemmer et al., 2016**). In other words, the test datasets—which include well-studied and characterised model systems— allow us to directly link the high AASD (as computed in this work) with high rates of sequence evolution (as found in previous works). This consideration can thus be extended also to the bivalve dataset: highly divergent genes in terms of AASD, which include some DSFGs and show an enrichment for GO-terms associated to macromolecule metabolism and morphological development (**Tab. 1**; **Supp. Tab. S8**), are also genes with accelerated substitution rates (**Ghiselli et al., 2018**; **Iannello et al., 2023**).

Concerning the second objection, we chose two test datasets with different characteristics as we wanted to check the extent of our hypothesis, that is, molecular evolution can be used to look for putative primary SDGs in taxonomic-wide analyses. The difference in divergence times and taxonomy ranks for bivalves and therians (Late Cambrian [about 498 million years ago, Mya; **Song et al., 2023**] and Early Mesozoic [166–123 Mya; **Álvarez-Carretero et al., 2022**], respectively) do not seem to influence the sequence diversity of SRGs, as both *Dmrt-1L*/*Sox-H* for bivalves and *Sry* for mammals exhibit high AASD with respect to their own distributions, regardless of their age. *Dmrt-1L* and *Sox-H* (which are mollusc- and Bilateria-specific, respectively) are undoubtedly older than *Sry,* which emerged in the Theria common ancestor (**Foster et al., 1992**), but each one of them can be considered a highly divergent gene in bivalves and mammals, respectively (i.e., genes that are included in the 5% upper quantile of bivalve and mammal AASD distributions). Conversely, the difference in divergence times and taxonomic ranks for *Drosophila* (Paleocene/Eocene boundary [about 56 Mya; **Russo et al., 2013**]) may seem to be influencing the results for the dataset, resulting in a false negative. In other words, it can be argued that: (i) the genes included in the SD cascade of *Drosophila* (such as *Sxl, tra*, and *dsx*; inset in **Fig. 4D**) may have a high AASD which, however, has not been detected by our methodological approach (for example, this may be traced back to the young diversification age of *Drosophila* species if compared to bivalves); (ii) the species included in the analysis are congeneric, thus the sequence differentiation of SRGs may exist not at the amino acid level but at the nucleotide one. To better disentangle this issue and further discuss the fruit fly dataset, we repeated the analysis of the AASD only on species of the *Crassostrea* genus (*C. gigas*, *Crassostrea angulata*, *Crassostrea ariakensis*, and *Crassostrea virginica*), which are much younger (Middle Cretaceous [less than 100 Mya; **Qi et al., 2023**]), thus comparable to *Drosophila*. Results showed that, even when analysing a smaller bivalve dataset, encompassing only 4 species of recent origin, the high AASD of *Dmrt-1L* persists, that is, *Dmrt-1L* is still grouped together with highly divergent genes (**Supp. Fig. S15**). The same has not been recovered for *Sox-H*, which fell in genes from Group 3 (the group corresponding to the 95% interval of the AASD distribution; see **Materials and Methods**) but still have the second highest AASD median value among DSFGs (**Supp. Fig. S15**).

Of course, we should not expect that highly divergent genes are only those involved in SD but may also participate in other processes (as discussed earlier and shown by GO term enrichments; **Tab. 1**; **Supp. Tab. S8**). Besides the genes of interest for SD (*Dmrt-1L*/*Sox-H* for bivalves, and *Sry* for mammals), other components of the DSFG families have been retrieved with a high AASD, despite they have never been linked directly to SD so far: *Sox-F* in bivalves (**Fig. 3B**) and *Fox-D4* in mammals (**Fig. 4B**). This implies that our approach cannot be used to unambiguously identify SDGs alone. Instead, the analysis is meant to be used to detect highly divergent genes and, subsequently, by comparison with literature and a more thorough and focused functional investigation, detect putative SDGs among them. In this view, the mammal dataset exemplifies the importance of putting the results of our pipeline (as those of any other comparative genomics analysis) into the correct evolutionary and genomic context: among DSFGs of mammals, two genes exhibit high AASD, one of which is directly related to SD (*Sry*), while the other has a function connected with neural development (*Fox-D4*; **Klein et al., 2013**). Thus, the high AASD may arise either because of the involvement in the upper SD pathway or because of other life-history traits connected with the gene, respectively. Regarding bivalves, *Dmrt-1L* and *Sox-H* show a sharp connection with SD as a putative primary SDG, either when considering their molecular evolutionary features or when looking at their gene expression and possible function in gonad development (**Naimi et al., 2009; Teaniniuraitemoana et al., 2014; Zhang et al., 2014; Capt et al, 2018; Li et al., 2018; Afonso et al., 2019; Liang et al., 2019; Yue et al., 2021**). It is difficult to further speculate on the actual involvement in SD of *Dmrt-1L* and *Sox-H* without any additional information on their biology, also considering that genes evolve not only at the level of their amino acid sequences, but also at the levels of exon-intron structures, indels, and regulatory relationships (among others). Nonetheless, molecular evolution proves to be a valuable tool to investigate genes putatively involved in SD, and to identify major targets onto which dedicate future research effort.

## CONCLUSIONS

Genes functioning in reproductive processes, and particularly SDGs, are often among the most variable in animal genomes, in terms of both sequence composition and regulatory interactions (**Swanson et al., 2001**; **Bachtrong et al., 2014**). Such high evolutionary rates may be traced back both to adaptive evolution (either as natural or sexual selection) or to non-adaptive processes (**Vicoso & Charlesworth, 2006**; **Meisel & Connallon, 2013**; **Parsch & Ellegren, 2013**; **Grath & Parsch, 2016**) and often results in striking differences in reproductive and sexual systems even among closely-related species. In the present work we took advantage of this characteristic to identify SDGs among the DSFG families in bivalves. By comprehensively analysing the phylogenetic history and AASD in a broad taxonomic dataset, we appointed *Dmrt-1L* and *Sox-H* as putative SDGs, thus confirming results in previous works that found them to be transcribed in a male-biased manner and/or strongly involved in male-gonad formation (**Naimi et al., 2009; Teaniniuraitemoana et al., 2014; Zhang et al., 2014; Capt et al, 2018; Li et al., 2018; Afonso et al., 2019; Liang et al., 2019; Yue et al., 2021**). Future studies would now need to further investigate their evolutionary history. For example, considering that SRGs tend to accumulate in the genomic neighbourhood where primary SDGs are located (**Capel, 2017**), analysing the genomic location of DSFGs in bivalve genomes may provide enlightening results. Similarly, revealing the genetic interactions of *Dmrt-1L* and *Sox-H*, through functional and genome editing assays, would undoubtedly benefit our understanding of their role in the sexual processes of bivalves.

## LIST OF ABBREVIATIONS

AASD: amino acid sequence divergence
DGE: differential gene expression
DM [domain]: *dsx* and *mab-3* [domain]
Dmrt: *doublesex* and *mab-3* related transcription factor
*Dmrt-1L*: *doublesex and mab-3 related transcription factor 1-like*
*Dm-W*: *Doublesex and mab-3 related gene W*
*Dmy*: *Doublesex and mab-3 related gene Y*
DSFGs: Dmrt, Sox, and Fox genes.
*dsx*: *doublesex*
Fox: forkhead box
ISH: *in-situ* hybridization
Mya: million years ago
SCO: single copy orthogroup
SD: sex determination
SDG: sex-determining gene
Sox: *Sry*-related HMG-box
SRG: sex-determination related gene
*Sry*: *Sex-determining region Y*

## ACKOWLEDGEMENTS

We would like to thank all the people who helped improving this work, by having promoted open discussions and/or exchanged ideas. But above all, we would like to thank Giobbe Forni, Mariangela Iannello and Giovanni Piccinini for always having offered their support whenever it was needed. This work was supported by the Canziani bequest funded to F.G. and A.L., by the European Union - NextGenerationEU under the National Recovery and Resilience Plan (PNRR) – Mission 4 Education and research – Component 2 From research to business - Investment 1.1, Notice Prin 2022 - DD N. 104 del 2/2/2022, entitled "Unraveling Sex Determination in Bivalves: An Integrative Single-Cell Analysis during the Early Embryo Development of *Mytilus galloprovincialis*", proposal code 2022CYM559 - CUP J53D23006200006 to L.M., and by the Saltonstall-Kennedy Program by NOAA Fisheries (SK-NOAA) funded to S.N.

## DATA ACCESSIBILITY AND BENEFIT-SHARING

### Data accessibility statement

The data underlying this article and the codes used to run the analyses are available on GitHub (https://github.com/filonico/bivalvia_SRGs).

### Benefit-sharing statement

Benefits from this research accrue from the sharing of our data, codes, and results on public databases as described above.

## AUTHOR CONTIRBUTIONS

FN, AL, FG, and LM conceived the study. FN performed research and analysed data. AL, FG and SN provided computational resources. FN wrote the paper. All authors edited and approved the manuscript.

## SUPPLEMENTARY TABLES

**Supp. Tab. S1. Genomic and transcriptomic data of bivalves and other molluscs.** For each species, the relative ID, taxonomic information, BUSCO statistics, NCBI accession number, and source publication are reported. Biv: Bivalvia; Ca: Caenogastropoda; Cep: Cephalopoda; Co: Coleoidea; Gas: Gastropoda; Gen: Genome; He: Heterobranchia; Im: Imparidentia; Ne: Neomphaliones; Pa: Palaeoheterodonta; Pt: Pteriomorpha; Tra: Transcriptome; Ve: Vetigastropoda.

**Supp. Tab. S2. Dmrt, Sox, and Fox gene (DSFG) family and domain identifiers (IDs) in PANTHER and CDD, respectively.** After having retrieved putative DSFGs based on HMM profiles, IDs have been used to retain only reliable hits.

**Supp. Tab. S3. List of DSFGs from reference species used to assess the identity of DSFGs in molluscs.** NCBI accession numbers are reported in parenthesis. Each row represents an orthology group.

**Supp. Tab. S4. Genomic data of mammals used to retrieve DSFGs and compute amino acid sequence divergence (AASD) of single copy orthogroups (SCOs).** For each species, the relative ID, taxonomic information, BUSCO statistics, NCBI accession number, and source publication are reported.

**Supp. Tab. S5. Genomic data of *Drosophila* used to retrieve DSFGs and compute AASD of SCOs.** For each species, the relative ID, taxonomic information, BUSCO statistics, NCBI accession number, and source publication are reported.

**Supp. Tab. S6. Complete set of DSFGs in bivalves.** For each gene, the species ID (Sp. ID) as in **Supp. Tab. S1**, the accession number (Gene ID), the Possvm-based annotation, and the CDD domains (including their Pssm-ID) are indicated.

**Supp. Tab. S7. Proportions of missing data in bivalve DSFGs.**

**Supp. Tab. S8. All the enriched GO terms for Group 1 and Group 2 genes of bivalves, mammals, and *Drosophila*.**

**Supp. Tab. S9. Complete set of DSFGs in mammals.** For each gene, the species ID (Sp. ID) as in **Supp. Tab. S4**, the accession number (Gene ID), and the Possvm-based annotation are indicated.

**Supp. Tab. S10. Complete set of DSFGs in *Drosophila*.** For each gene, the species ID (Sp. ID) as in **Supp. Tab. S5**, the accession number (Gene ID), and the Possvm-based annotation are indicated.

## SUPPLEMENTARY FIGURES

**Supp. Fig. S1. ML phylogenetic tree of the Dmrt gene family in molluscs, including the Possvm orthology inference.** Reference genes from *H. sapiens*, *C. elegans*, and *D. melanogaster* are marked with an asterisk at the beginning of the tip. Species ID can be found in **Supp. Tab. S1**. Bootstrap values are shown for each node.

**Supp. Fig. S2. ML phylogenetic tree of the Sox gene family in molluscs, including the Possvm orthology inference.** Reference genes from *H. sapiens*, *C. elegans*, and *D. melanogaster* are marked with an asterisk at the beginning of the tip. Species ID can be found in **Supp. Tab. S1**. Bootstrap values are shown for each node.

**Supp. Fig. S3. ML phylogenetic tree of the Fox gene family in molluscs, including the Possvm orthology inference.** Reference genes from *H. sapiens*, *C. elegans*, and *D. melanogaster* are marked with an asterisk at the beginning of the tip. Species ID can be found in **Supp. Tab. S1**. Bootstrap values are shown for each node.

**Supp. Fig. S4. Dmrt, Sox, and Fox gene (DSFG) complement in Mammalia and *Drosophila* spp.** Presence/absence of genes in various species are indicated by filled circles. Numbers inside each circle specify genes with 2 or more copies. The shaded area highlights outgroup species, *Gallus gallus* (Aves) for mammals and *Anopheles gambiae* (Culicidae) for fruit flies. The phylogenetic tree of analyzed species, as inferred from literature, is shown on the left, while major taxonomic groups are reported on the right. All species are represented by genomic data. DSFG trees are shown on the bottom (full trees can be found in **Supp. Fig. S5–S7**). Full species names for both mammals and fruit flies, along with all assembly and taxonomic information, can be found in **Supp. Tab. S4** and **Supp. Tab. S5**, respectively. A.: Aves; Chirop.: Chiroptera; L.: Lagomorpha; M.: Monotremata; Me.: Metatheria; P.: Pholidota; Pe.: Perissodactyla; Prim.: Primates; Roden.: Rodentia; X.: Xenarthra; C.: Culicidae.

**Supp. Fig. S5. ML phylogenetic tree of the Dmrt gene family in mammals, including the Possvm orthology inference.** Reference genes from *H. sapiens*, *Mus musculus*, *Elephas maximus indicus*, and *Ornithorhynchus anatinus* are marked with an asterisk at the beginning of the tip. Species ID can be found in **Supp. Tab. S4**. Bootstrap values are shown for each node.

**Supp. Fig. S6. ML phylogenetic tree of the Sox gene family in mammals, including the Possvm orthology inference.** Reference genes from *H. sapiens*, *Mus musculus*, *Elephas maximus indicus*, and *Ornithorhynchus anatinus* are marked with an asterisk at the beginning of the tip. Species ID can be found in **Supp. Tab. S4**. Bootstrap values are shown for each node.

**Supp. Fig. S7. ML phylogenetic tree of the Fox gene family in mammals, including the Possvm orthology inference.** Reference genes from *H. sapiens*, *Mus musculus*, *Elephas maximus indicus*, and *Ornithorhynchus anatinus* are marked with an asterisk at the beginning of the tip. Species ID can be found in **Supp. Tab. S4**. Bootstrap values are shown for each node.

**Supp. Fig. S8. ML phylogenetic tree of the Dmrt gene family in fruit flies, including the Possvm orthology inference.** Reference genes from *Drosophila melanogaster*, *Drosophila hydei*, *Drosophila pseudoobscura*, and *Drosophila suzukii* are marked with an asterisk at the beginning of the tip. Species ID can be found in **Supp. Tab. S5**. Bootstrap values are shown for each node.

**Supp. Fig. S9. ML phylogenetic tree of the Sox gene family in fruit flies, including the Possvm orthology inference.** Reference genes from *Drosophila melanogaster*, *Drosophila hydei*, *Drosophila pseudoobscura*, and *Drosophila suzukii* are marked with an asterisk at the beginning of the tip. Species ID can be found in **Supp. Tab. S5**. Bootstrap values are shown for each node.

**Supp. Fig. S10. ML phylogenetic tree of the Fox gene family in fruit flies, including the Possvm orthology inference.** Reference genes from *Drosophila melanogaster*, *Drosophila hydei*, *Drosophila pseudoobscura*, and *Drosophila suzukii* are marked with an asterisk at the beginning of the tip. Species ID can be found in **Supp. Tab. S5**. Bootstrap values are shown for each node.

**Supp. Fig. S11. ML phylogenetic tree of the Dmrt gene family in mollusc species.** Species ID can be found in **Supp. Tab. S1**. The tree has been midpoint rooted. Bootstrap values are shown for each node.

**Supp. Fig. S12. ML phylogenetic tree of *Sox-B1* and *Sox-B2* genes in mollusc and reference species**. Reference genes from *H. sapiens*, *C. elegans*, and *D. melanogaster* are marked with an asterisk at the beginning of the tip names. Species ID can be found in **Supp. Tab. S1**. Bootstrap values are shown for each node.

**Supp. Fig. S13. ML phylogenetic tree of *Fox-J2*, *Fox-M*, *Fox-O*, and *Fox-P* genes in mollusc and reference species.** Reference genes from *H. sapiens*, *C. elegans*, and *D. melanogaster* are marked with an asterisk at the beginning of the tip names. Species ID can be found in **Supp. Tab. S1**. Bootstrap values are shown for each node.

**Supp. Fig. S14. ML phylogenetic tree of the Fox gene family in bivalves and the sea urchin *Strongylocentrotus purpuratus* (Spur).** Reference genes from *S. purpuratus* are marked with an asterisk at the beginning of the tip names. Species ID can be found in **Supp. Tab. S1**. *S. purpuratus* genes are those given by **Tu et al., 2006**. Bootstrap values are shown for each node.

**Supp. Fig. S15. Distribution of amino acid sequence divergence (AASD) of single-copy orthogroups in *Crassostrea gigas*, *Crassostrea angulata*, *Crassostrea ariakensis*, and *Crassostrea virginica* (A), including DSFG (B)**. The distribution of AASD in *Crassostrea* has been computed on the median values of pairwise distances of over 14 k single-copy orthogroups (SCOs). Circle heights of DSFGs show the median value of their AASD. *Dmrt-1L* genes are indicated as ‘dmrt_disco_tree3’.

